# A comparison of four proteomics software for hair proteome analyses – A technical note

**DOI:** 10.64898/2026.04.17.719199

**Authors:** M. Mukonyora

**Affiliations:** Department of Medicine, University of Cape Town, Groote Schuur Hospital, Cape Town, South Africa

## Abstract

Hair has applications in biomarker discovery and forensics, yet the influence of proteomics software tools on hair proteome characterisation remains underexplored. This study compares four bottom-up proteomics workflows (MaxQuant, FragPipe, MetaMorpheus, and SearchGUI/PeptideShaker). Publicly available hair proteomes were analysed following extraction with 1-dodecyl-3-methylimidazolium chloride (DMC), sodium dodecanoate (SDD), sodium dodecyl sulfate (SDS), and urea. Data were acquired on Orbitrap-based DDA platforms. Peptide identification, protein inference, functional annotation, physicochemical properties, and label-free quantification (LFQ) were evaluated. Peptide-level performance differed across tools. MS-GF+ and FragPipe identified the most unique peptides, while X!Tandem reported the fewest. Protein inference showed a dissociation from peptide-level results. MetaMorpheus reported the highest number of protein groups despite only the third highest peptide counts. FragPipe and MaxQuant followed, while PeptideShaker consistently inferred the fewest proteins. Protein-level concordance was low, with only 30.3% overlap across tools and extraction methods. These differences extended to downstream analyses. Functional enrichment showed moderate concordance (38.25% overlap). Physicochemical profiles varied, with MetaMorpheus identifying more hydrophobic proteomes and PeptideShaker more hydrophilic profiles. At the quantitative level, reproducibility depended on extraction buffer. SDS and urea showed lower variability (CV =< 0.025), while DMC and SDD showed higher variability (up to 0.10). Absolute LFQ intensities and differential expression outputs varied across tools despite moderate to strong correlation (r = 0.77 to 0.93). Overall, software choice influences proteome coverage, physicochemical profiles, and quantitative outcomes. Relative trends were partially conserved, but magnitude and significance varied. These findings support careful method selection and multi-tool validation in hair proteomics

## 1.2 Introduction

Hair proteomics is an emerging field yet no study to date has systematically evaluated the impact of software choice on the characterisation of the hair proteome. The hair proteome harbours valuable disease biomarkers, protein profiles associated with hair curvature and cosmetic properties, as well as single amino acid polymorphisms (SAPs) applied in forensics [1-5]. During keratinisation, a terminal differentiation programme, the hair shaft becomes highly abundant in keratins while some cellular components are degraded, and others like ribosomes and mitochondria are maintained to drive the process [6, 7]. The result is an increased dynamic range, which complicates the identification of less abundant proteins that may be of interest as disease biomarkers [8]. Hair’s resilient structure, reinforced by extensive disulfide and ε(γ–glutamyl) lysine isopeptide bond networks, presents significant challenges for protein extraction [9-11]. Extraction protocols are classified into detergent- and non-detergent-based buffers, each affecting the complement of recovered proteins [9-13]. However, there is no consensus on the optimal extraction method for hair proteins.

Mass spectrometry, particularly Orbitrap-based data-dependent acquisition (DDA), is the most widely used approach for hair proteome analyses [4, 10, 11, 14-16]. In bottom-up proteomics, peptide identification relies on with either de novo sequencing or database searching. De novo methods utilise fragmentation rules and probabilistic models (e.g., Hidden Markov Models and Graphical Probabilistic Models), enabling detection of variants absent from reference databases. However, they are constrained by incomplete fragmentation and lower mass accuracy [17-19]. In contrast, database search engines such as MASCOT [20], Andromeda [21], SEQUEST [22], and X!Tandem [23] match spectra to predicted peptides using scoring algorithms and user-defined parameters [24]. Following identification, protein inference reconstructs protein composition using approaches like the parsimonious rule, hierarchical statistical models, and Bayesian inference. The accuracy of protein inference depends on robust peptide-spectrum match (PSM) scores and stringent cutoffs [17, 25].

Proteomics quantification may either be absolute or relative, with or without the use of labels [26, 27]. Relative label-free quantification (LFQ) is widely applied in hair proteomics studies [3, 4, 10, 11, 14, 15, 28]. It comprises multiple steps, including data transformation, normalization, and imputation [27, 29, 30]. Relative LFQ primarily relies on either peak intensity-based methods or spectral counting, the former of which is typically more accurate due to its use of MS1 signal intensities and feature alignment strategies [27, 31].

Proteomics software pipelines may be also grouped into modular workflows that combine independent tools, and non-modular (integrated) workflows, which handle the entire process within a single platform. Both commercial and open-source solutions are available [26, 29]. These bioinformatics workflows can be complex and unintuitive, particularly for novice users without computational backgrounds, thereby limiting accessibility to non-specialists. To address these challenges, different GUI-based software suites offering protein inference and diverse quantification strategies were developed for users with varying expertise levels [32]. MaxQuant is a widely adopted non-modular software incorporating the Andromeda search engine [33, 34]. FragPipe is a modular pipeline built around the MSFragger search engine notable fast for its computational efficiency on standard laptops [35]. It integrates tools including Philosopher and IonQuant for protein inference and quantification, providing flexibility and scalability [36, 37]. MetaMorpheus is a user-friendly, non-modular platform that can run large-scale search tasks on memory-restricted machines [38]. It also offers flexible data analysis workflows for users’ specific needs [39]. PeptideShaker offers a modular, user-friendly framework for protein inference. It is compatible with SearchGUI, which leverages multiple search engines including Andromeda, Comet, Mascot, MS Amanda, SequestHT, and X!Tandem [32, 40, 41]. Protein inference software is an important consideration because it impacts in the results of the research question, the quantitation values and the final claims in the research manuscript.

Given the complexity of extracting hair proteins and challenges identifying less abundant proteins due to the high abundance of keratins, we reviewed the literature to compare existing hair extraction buffers and proteomics software pipelines. We then evaluated four widely used bottom-up proteomics pipelines (modular and non-modular) to investigate their impact on hair protein identification and LFQ. Namely, MaxQuant (Andromeda), FragPipe (MSFragger), MetaMorpheus, and SearchGUI (five search engines) coupled with PeptideShaker. Our findings provide guidance toward the selection of proteomics software pipelines for hair proteomics, particularly for disease biomarker discovery and forensic applications.

## 1.3 Methods

We assessed the impact of search engine selection on hair peptide identification and protein inference. We selected freely accessible search engines compatible with DDA data, which can be run on a laptop with at least 16GB of RAM: FragPipe, MaxQuant, MetaMorpheus, and SearchGUI/PeptideShaker. Within SearchGUI *v4*.*3*.*11* we further compared five search engines, namely Andromeda, X!Tandem, OMSSA, MSAmanda, and MSGF+. A literature search was conducted to identify hair proteomics studies with publicly available MS data, with an emphasis on selecting different protein extraction buffers.

Four hair proteomes were therefore downloaded from ProteomeXChange, whose extractions used 1-dodecyl-3-methylimidazolium chloride (DMC), sodium dodecyl sulfate (SDS), sodium dodecanoate (SDD), and urea respectively. Additionally, only datasets generated using Orbitrap mass spectrometers operating in DDA mode were considered. The ProteomeXChange IDs are PXD007224 (urea and SDS) [11], PXD011732 (SDD) [14], and PXD034814 (DMC) [4]. We used three biological replicates for each extraction method, except for the SDS and urea methods where only two replicates were available (**Table 1**).

**Table 1:** Four hair proteomic datasets used in this study. These studies used extraction buffers with1-dodecyl-3-methylimidazolium chloride (DMC), sodium dodecyl sulfate (SDS), sodium dodecanoate (SDD), and urea respectively. Their mass spectrometry (MS) data was generated using Orbitrap mass spectrometers in DDA mode, and the raw MS data was publicly available.

| Citation | Detergent/denaturant | Mass or length of hair | Number of samples used in this study | Homogenization | Hair classification | Analytical platform | DDA or DIA | Proteome XChange ID |
| --- | --- | --- | --- | --- | --- | --- | --- | --- |
| Adav *et al.* 2018. [11] | urea | 50mg | 2 (no fractions) | Not specified | Unknown | Nano-LC coupled to a Q-Exactive (Orbitrap-Q-ToF) | DDA | PXD007224 |
| Adav *et al.* 2018. [11] | sodium dodecyl sulfate | 80mg | 2 (2 fractions per sample) | Not specified | Unknown | Nano-LC coupled to a Q-Exactive (Orbitrap-Q-ToF) | DDA | PXD007224 |
| Milan *et al.* 2019 [11] | sodium dodecanoate | 25mg and 2 inches long | 3 (no fractions) | Cut 2mm | Unknown | Nano-LC coupled to a Q-Exactive Orbitrap | DDA | PXD011732 |
| Wu *et al.* 2022 [4] | 1-dodecyl-3-methylimidazolium chloride | 2cm single hair | 3 (6 fractions per sample) | cut 1–2-mm | Han Chinese | 2D Nano-LC coupled to an Orbitrap Fusion Lumos Tribrid | DDA | PXD034814 |

Protein inference was carried out using a target and decoy version of the UniProt reference human proteome FASTA file (Proteome ID: UP000005640; accessed June 2021). Search parameters and post-processing settings were standardized across pipelines. The fixed modification was carbamidomethylation of cysteine. Five variable modifications were used of N-terminal acetylation, oxidation of methionine, deamidation of N, deamidation of Q, and ubiquitination of K. Up to two missed cleavages of trypsin were allowed. The precursor tolerance was set as 10ppm, and the fragment tolerance was up to 0.02 Da. The peptide and protein false discovery rate (FDR) was 0.01, and the default minimum peptide length of 8 amino acids was allowed. The fragment ion types were set as b and y. LFQ using peak intensity was conducted using match between runs and default settings for FragPipe, MaxQuant, and MetaMorpheus. PeptideShaker was excluded from the quantification comparison because it uses spectral counts.

The unique and shared proteins among the different protein inference algorithms were determined using a free online Venn diagram plotting software called Venny [42]. To qualitatively investigate the impact of algorithm choice and the type of extraction method on the proteins identified, gene ontology (GO) enrichment analysis was performed with the online software STRING *v11*.*5* [43]. We compared the GO analyses of all proteins identified by each software to determine whether their inferred proteins deliver consistent biological insights. All analyses were conducted using R software *v2025*.*05*.*1+513*. To characterize the hydrophobicity patterns of proteomes identified by the respective software platforms we calculated their distribution of Grand Average of Hydropathicity (GRAVY) scores.

To ensure robust quantitative comparisons, we compared the protein abundance profiles using two or three biological replicates per extraction method across the three proteomics software tools. Proteins detected across all tools for each extraction method were retained for quantitative comparison. These shared proteins were organized into matrices with proteins as rows and samples as columns using an in-house R-script. Their LFQ data were log2 transformed and missing values were imputed using a normal distribution in the Perseus software *v2*.*1*.*5*.*0*. For each extraction method, R was used to calculate Pearson correlations, distributions, coefficients of variation (CV), and differential expression analyses of the respective tools.

## 1.4 Results and Discussion

### 1.4.1 Peptide Identification Sensitivity Across Search Engines

Quality assessment of peptide sequencing and protein inference showed substantial variation in identification performance across software platforms (FragPipe, MaxQuant, MetaMorpheus, SearchGUI/PeptideShaker). MS-GF+ within the PeptideShaker framework achieved the highest average sequence coverage of the hair proteome (26.3%), while MetaMorpheus reported the lowest (13.55%) across all extraction buffers (DMC, SDD, SDS, urea) (**Figure 1a**). This nearly two-fold difference underscores divergence in peptide detection sensitivity between tools.

**Figure 1.**
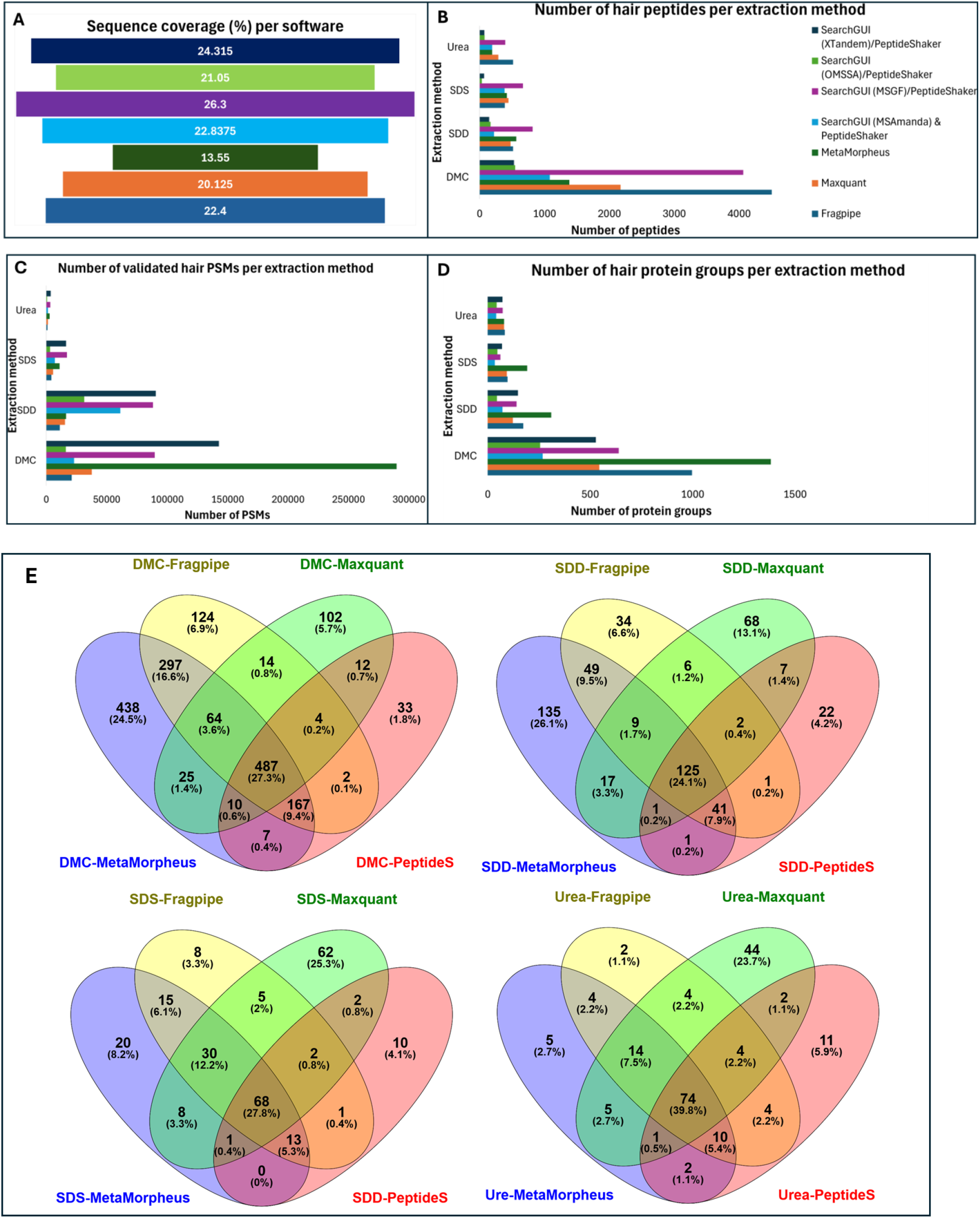
Peptide and protein identification across software tools and extraction methods. (a) Average sequence coverage (%) of the hair proteome across search engines for each extraction method (DMC, SDD, SDS, urea). (b) Number of peptide-spectrum matches (PSMs). (c) Number of validated hair PSMs. (d) Number of unique hair peptides identified per software tool. (e) Overlap and unique protein groups inferred across software tools and extraction methods.

At the PSM level, MS-GF+ and X!Tandem identified the greatest numbers of PSMs, whereas OMSSA consistently identified the fewest (**Figure 1b**). In contrast, unique peptide identification followed a different trend, with MS-GF+ and FragPipe outperforming other tools, while X!Tandem reported the lowest unique peptide counts across three out of four extraction buffers (**Figure 1c; Supplementary Table 1**). This divergence highlights that higher PSM counts do not necessarily translate into improved proteome coverage but may instead reflect redundant matching of spectra to the same peptide sequences [24]. The strong performance of MS-GF+ is consistent with its probabilistic scoring model [44]. FragPipe’s performance likely reflects MSFragger’s ability to efficiently search expanded peptide spaces, including fragment ion indexing and open search strategies, which have been reported to increase PSMs by up to 32% [35, 36]. These findings are supported by recent benchmarking studies showing improved peptide-level performance of MSFragger-based workflows relative to MetaMorpheus and MaxQuant [45, 46]. Conversely, X!Tandem’s lower unique peptide yield despite high PSM counts suggests repeated identification of spectra derived from abundant peptides, consistent with its scoring approach [47]. OMSSA’s lower PSM counts likely reflect a more conservative scoring strategy that rejects lower-confidence spectra [48].

### 1.4.2 Protein Inference and Unique Protein Recovery

Protein inference revealed a dissociation between peptide- and protein-level performance. MetaMorpheus reported the third highest number of unique peptides but the highest number of protein groups across extraction methods, followed by FragPipe and MaxQuant, while PeptideShaker inferred the fewest. Similarly, within the SearchGUI/PeptideShaker suite, X!Tandem inferred the largest number of protein groups across three of four extraction buffers (SDD, SDS, urea), despite identifying the fewest unique peptides (**Figure 1d**). This apparent paradox reflects ambiguity in peptide-to-protein mapping, where shared peptides reduce the specificity of protein inference. Differences between tools arise from how this ambiguity is resolved. MetaMorpheus likely reports more protein groups due to less aggressive collapsing of protein variants prior to grouping, whereas MaxQuant and PeptideShaker implement stricter parsimony-driven grouping strategies that reduce protein group counts [38, 49]. FragPipe and MaxQuant both balance parsimony and protein coverage through heuristic approaches; however, FragPipe reports more protein groups, potentially reflecting differences in protein inference and grouping within the ProteinProphet framework [24]. PeptideShaker applies parsimony-based protein inference, in which shared peptides introduce ambiguity and promote the formation of indistinguishable protein groups. However, its overall implementation favours more conservative protein reporting, resulting in fewer inferred protein groups [40, 50].

Protein-level concordance between software tools was low, underscoring the impact of search and inference strategies on the resulting proteome. Only 30.3% of proteins were shared across all four tools and extraction methods (**Figure 1e**), lower than the ∼55% overlap reported in plasma/serum studies [51]. MetaMorpheus reported the highest unique protein proportions for DMC (24.5%) and SDD (26.1%), while MaxQuant led for SDS (25.5%) and urea (23.7%). FragPipe yielded the lowest unique proportions for SDS (3.3%) and urea (1.1%), and PeptideShaker the lowest for DMC (1.8%) and SDD (4.2%) (**Figure 1e**). These differences reflect variation in proteome search space expansion and inference stringency. FragPipe achieved a balance between moderate protein counts and low unique proportions, suggesting a more conservative and robust identification strategy.

Because all datasets were acquired on Orbitrap platforms and processed using identical search parameters, differences in protein identifications are unlikely to arise from instrument variability or parameter inconsistencies. Instead, variation in unique protein proportions reflects the combined effects of extraction chemistry, biological heterogeneity of hair, and software-specific scoring and protein inference frameworks [24, 40, 52]. Even under harmonized conditions, search engines interpret identical spectral evidence using distinct statistical models, leading to divergent protein-level outputs. Establishing a reference hair proteome or ground-truth dataset will be critical for disentangling true biological variability from software-driven differences, and for benchmarking protein inference performance across tools [53].

### 1.4.3 Functional Enrichment Consistency

Functional enrichment analysis showed moderate concordance across software tools, with an average overlap of 38.25%. Reactome pathway predictions were most consistent (54.9%), while GO biological process and molecular function terms showed lower agreement (28.4% and 33.5%) (**Figure 2**). This is consistent with differences in protein inference and identification outputs across proteomics workflows and the propagation of these differences into downstream functional annotation [27, 40]. FragPipe produced the highest number of unique functional terms, indicating broader proteome coverage, whereas PeptideShaker yielded the fewest, consistent with more conservative protein inference (**Figure 2**). These findings highlight that pathway-level conclusions in biomarker discovery can be workflow-dependent, underscoring the need for standardized benchmarking of proteomics pipelines prior to biological interpretation [24, 54].

**Figure 2.**
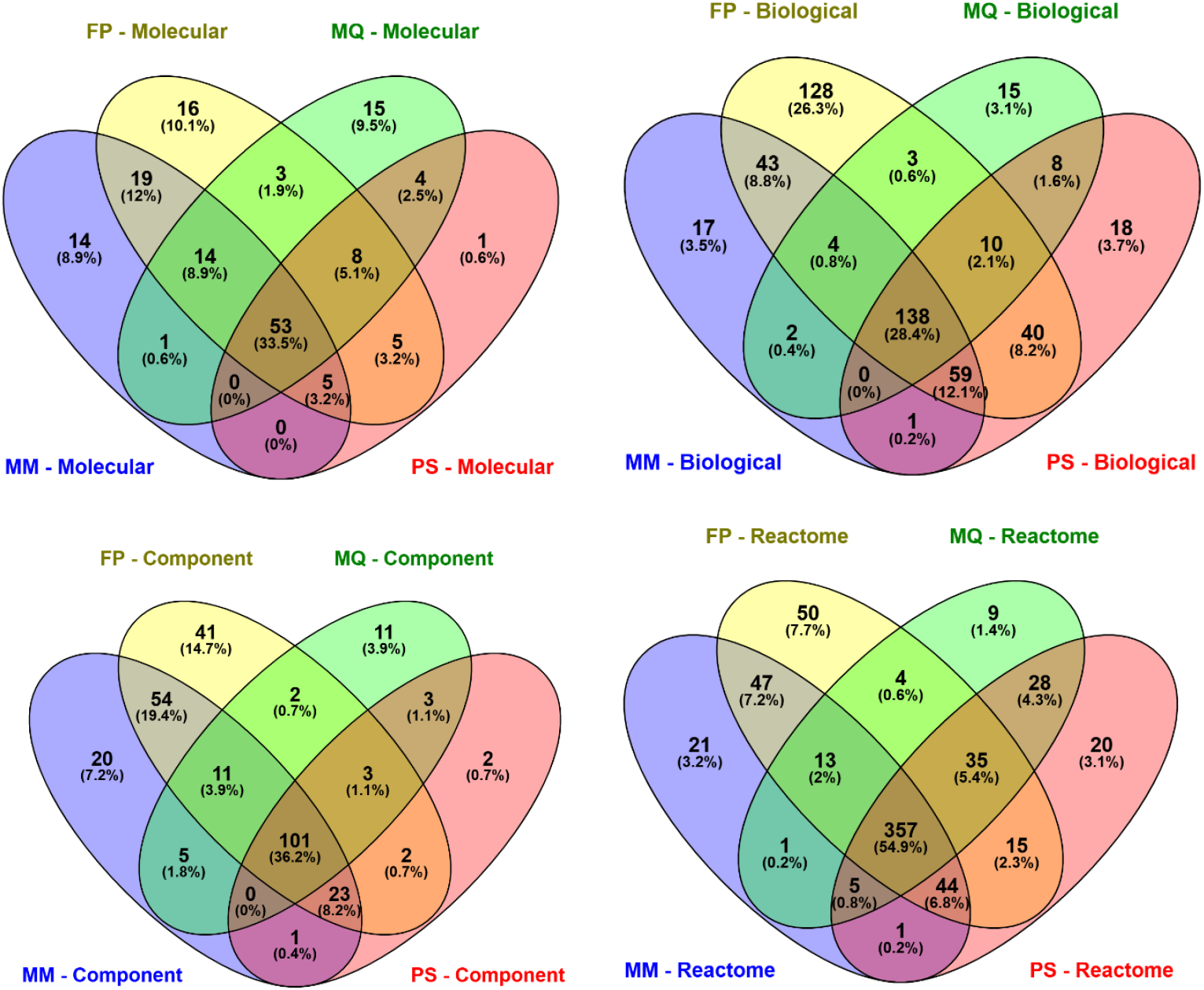
Functional enrichment overlap across software-derived protein lists. Overlap of Gene Ontology (molecular function, biological process, cellular component) and Reactome pathway enrichment results across software tools. Values represent percentage overlap between tool-derived protein sets.

### 1.4.4 Hydrophobicity Profile (GRAVY)

GRAVY scores were calculated to assess the hydrophobicity of proteins identified by each software tool. The overall distribution shapes were comparable across tools indicating no major systematic bias in hydrophobicity profiling. However, GRAVY ranges varied, with MetaMorpheus (-1.67 to 0.79) identifying the most hydrophobic proteins (highest GRAVY) and PeptideShaker (-1.88 to 0.49) the most hydrophilic (lowest GRAVY). MaxQuant (-1.29 to 0.15) and FragPipe (-1.28 to 0.58) showed narrower ranges (**Figure 3**).

**Figure 3.**
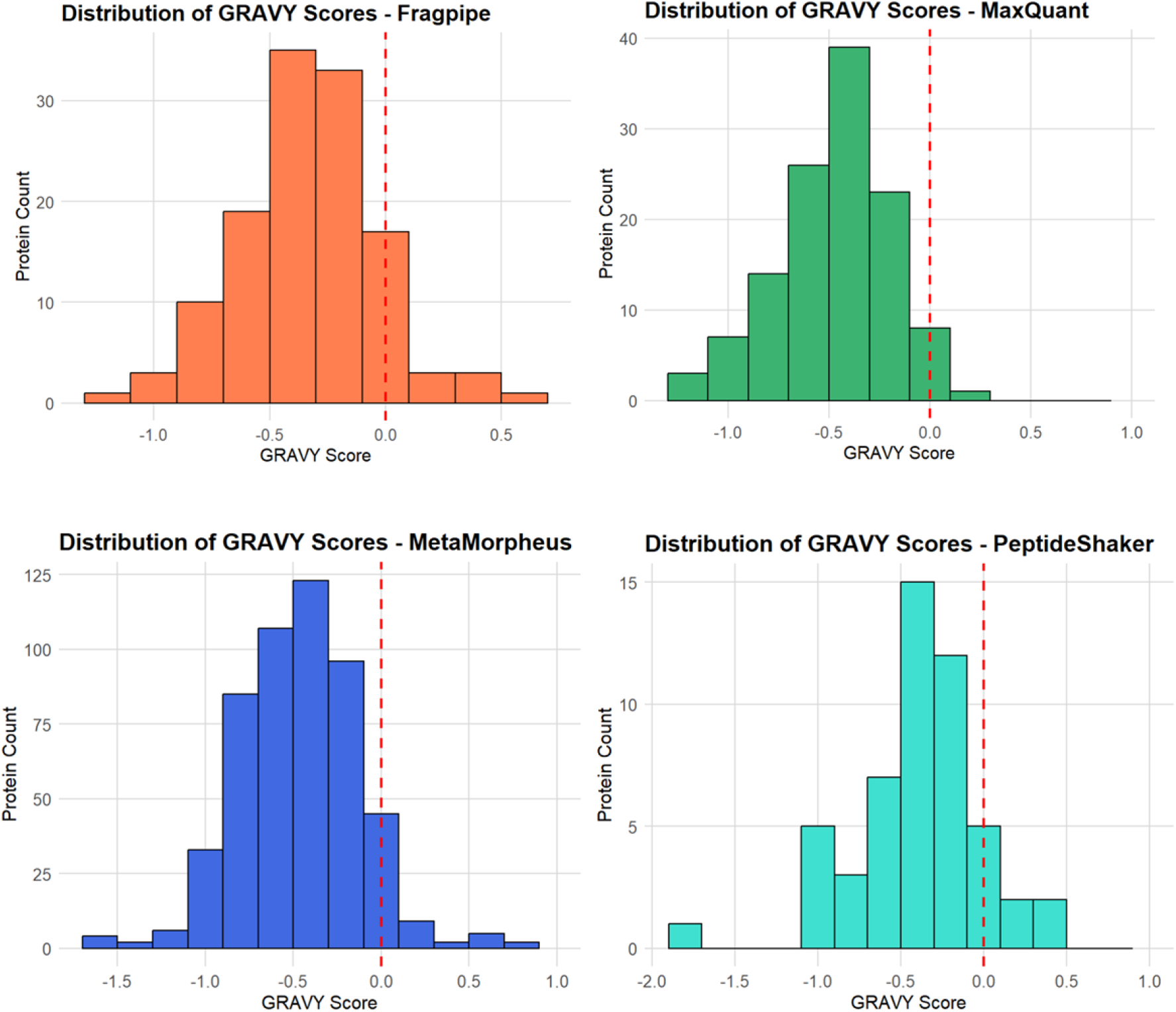
GRAVY score distributions of identified proteins. Distribution of GRAVY hydrophobicity scores for proteins identified by each software tool across all extraction methods.

Differences in peptide identification and protein inference strategies across search engines determine the set of peptides and proteins reported, which in turn influences downstream physicochemical characterisation of the proteome. In LC–MS-based LFQ workflows, peptide detectability is inherently biased by hydrophobicity-dependent processes, including ionization efficiency, chromatographic retention, and fragmentation behaviour [27, 55]. As a result, the observed hydrophobicity distribution of identified proteins reflects both experimental selection effects and software-specific decision rules, particularly in complex matrices such as hair [24, 40]. The small hydrophobic KAPs are often lost along the proteomic pipeline from extraction, tryptic digestion, to LC-MS analysis [56-59].

### 1.4.5 Quantitative Reproducibility and Agreement

To assess reproducibility of the LFQ, the CVs of the respective software were compared for each extraction buffer. SDS and urea provided higher quantitative reproducibility (CV ≤ 0.025) compared to DMC and SDD (up to 0.10) (**Figure 4**). This indicates lower variability in protein-level quantification under SDS and urea conditions within the LFQ proteomics workflows [27]. Keratins and KAPs were prominent among the top 10 most variable proteins for DMC (6 KAPs), urea (5 KAPs, 2 keratins), and SDS (1 keratin, 4 KAPs), reflecting the known challenges of extracting highly crosslinked hair proteins (**Supplementary Tables 2-5**) [9, 60]. In contrast, SDD variability was dominated by non-structural proteins (2 KAPs), indicating preferential extraction of the more soluble hair protein fractions (**Supplementary Tables 2-5**). Overall, these findings suggest inconsistent recovery of highly crosslinked and insoluble hair structural proteins by the different extraction buffers, suggesting a trade-off between reproducibility and proteome accessibility [61, 62].

**Figure 4.**
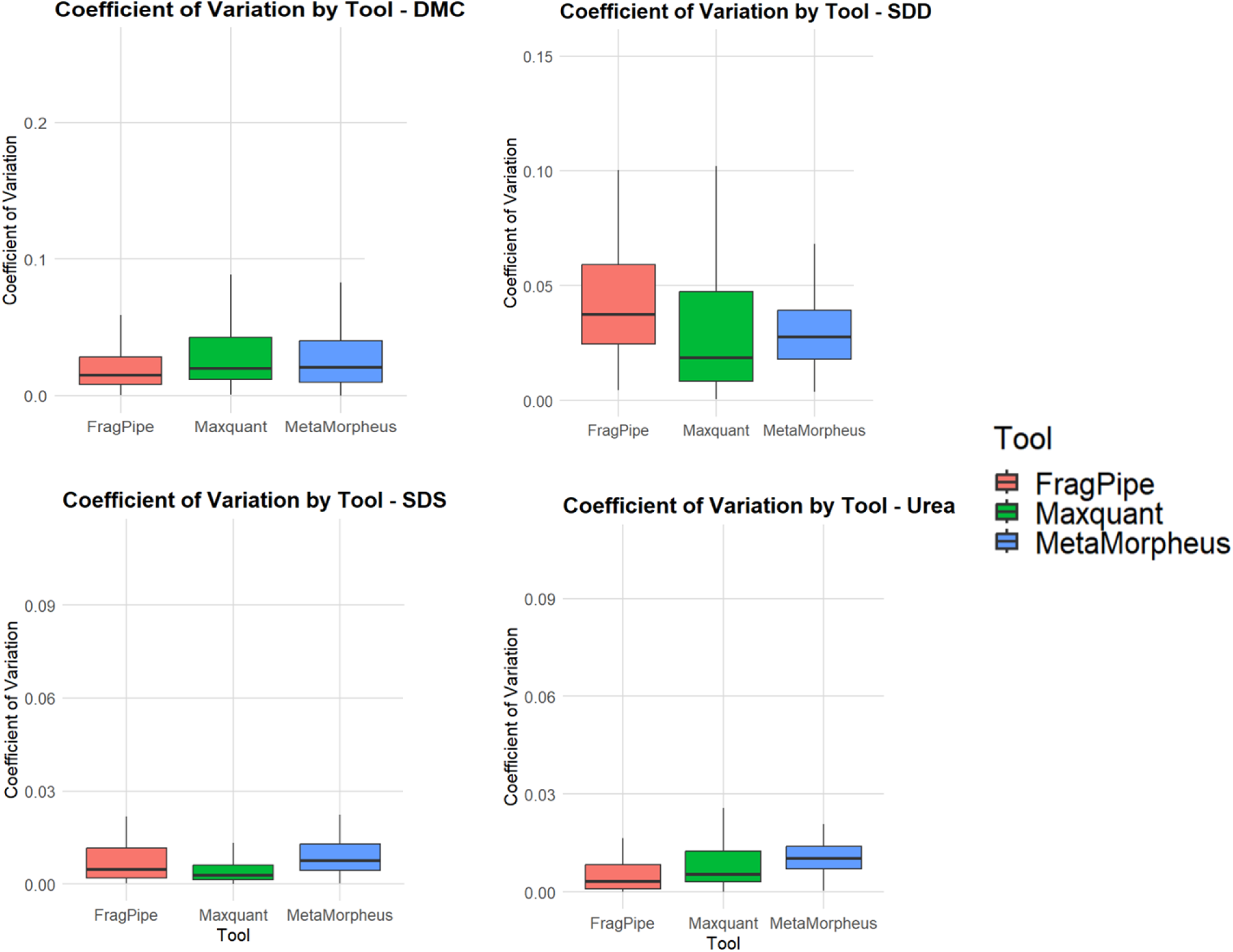
Coefficient of variation across extraction methods. Protein-level coefficients of variation (CV) of LFQ intensities across extraction methods (DMC, SDD, SDS, urea) for each software tool.

Boxplots of LFQ intensity distributions across tools showed that MetaMorpheus consistently reported the lowest median values, while MaxQuant yielded the highest median intensities for all extraction buffers except urea (**Figure 5**). These systematic shifts in intensity distributions are consistent with previously reported inter-software variability in LFQ workflows and likely arise from differences in normalization strategies, protein inference, and intensity aggregation methods [26, 34, 55, 63].

**Figure 5.**
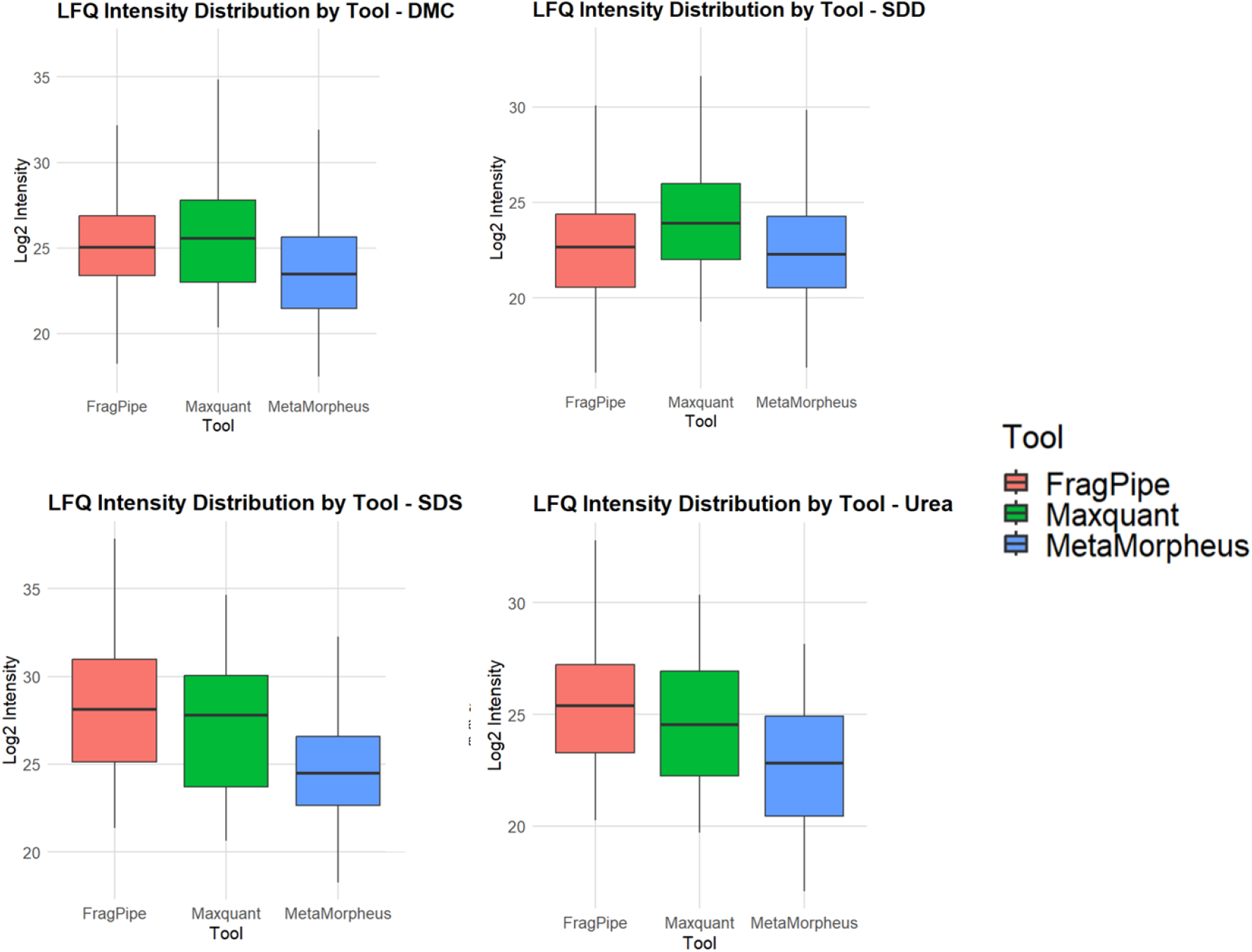
LFQ intensity distributions across software platforms. Boxplots showing LFQ intensity distributions for each software tool across all extraction methods.

We assessed quantitative agreement by calculating Pearson correlation coefficients between protein inference tools. Moderate to strong quantitative concordance was observed, with the degree of agreement varying by extraction chemistry. Pearson correlations showed moderate to strong agreement (r = 0.77–0.93), with strongest concordance for SDD (FragPipe vs MetaMorpheus, r = 0.93) and lower agreement for urea (0.77–0.84) (**Figure 6**).

**Figure 6.**
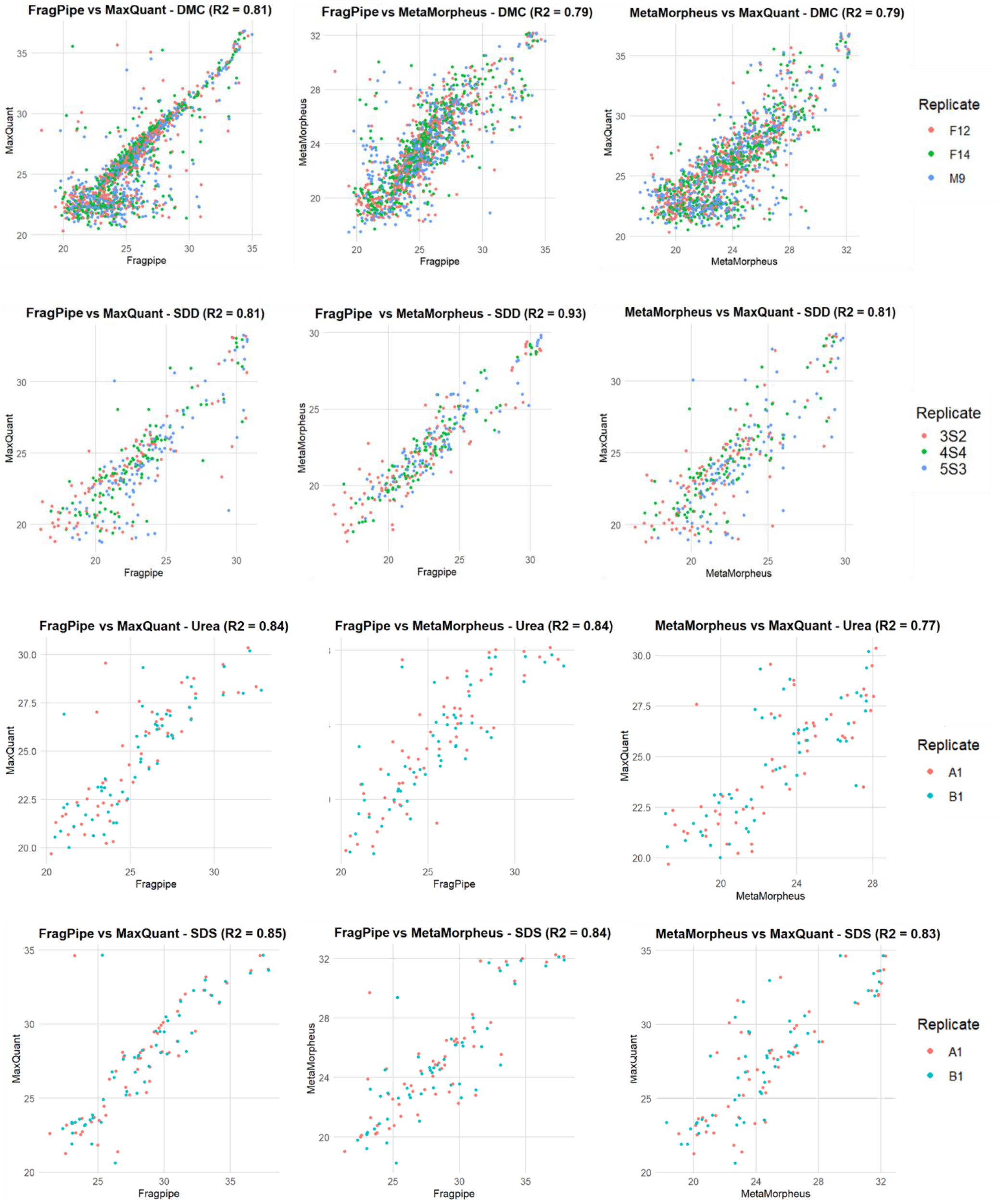
Pairwise correlation of LFQ intensities between software tools. Pearson correlation coefficients of protein-level LFQ intensities between software tools for each extraction method.

Differential expression analysis using normalized LFQ values confirmed systematic differences in quantitative output across software tools, with MetaMorpheus reporting significantly higher protein intensities relative to both FragPipe and MaxQuant (p < 0.05). Consequently, MetaMorpheus identified a greater number of upregulated proteins, whereas FragPipe and MaxQuant reported few downregulated proteins in pairwise comparisons. The only instance of substantial downregulation was observed for FragPipe relative to MaxQuant (**Figure 7**). These results indicate that software-dependent intensity scaling and protein inference can influence downstream differential abundance results, affecting biological interpretations [34, 63].

**Figure 7.**
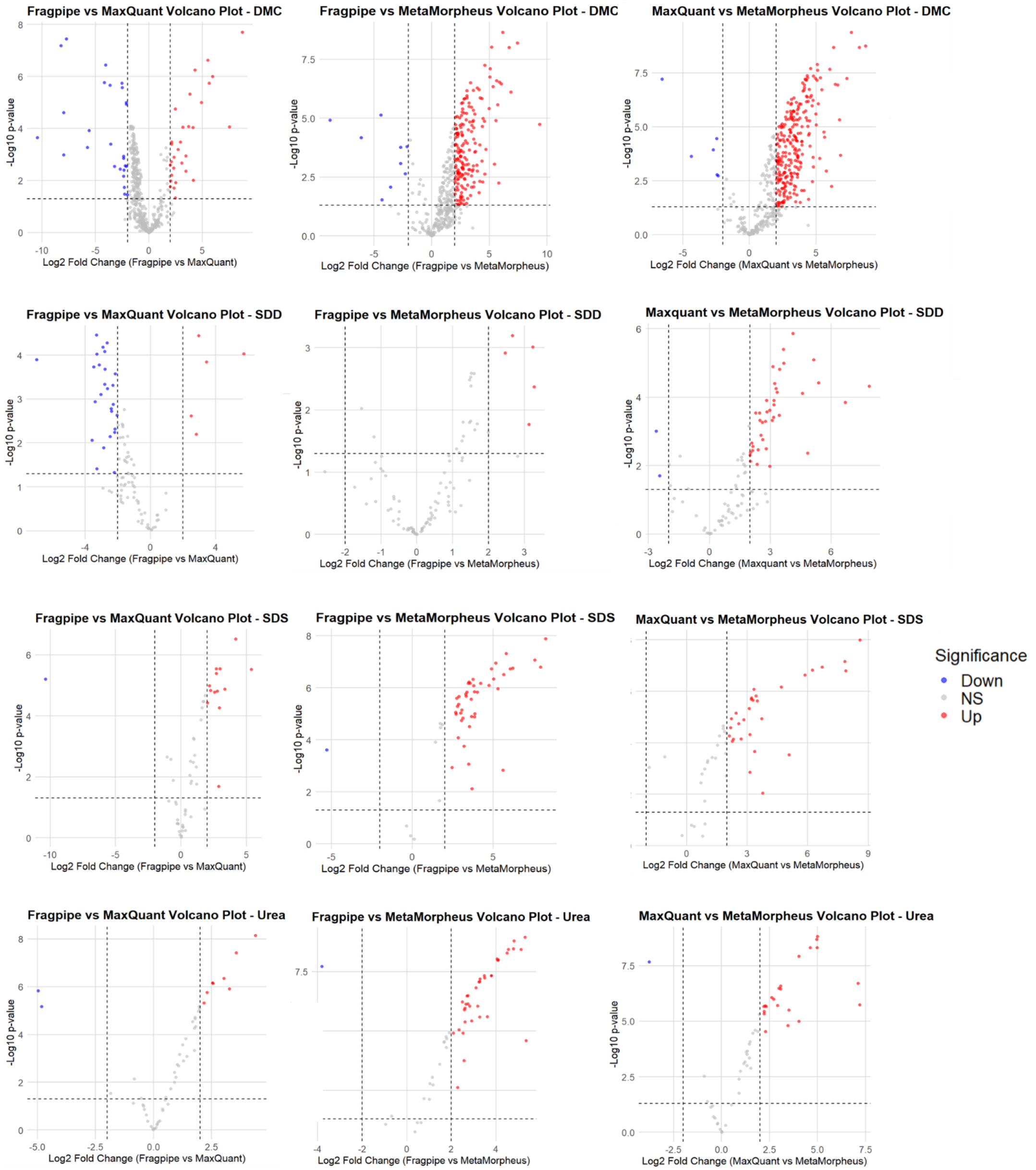
Differential expression across software tools. Number of significantly up- and downregulated proteins identified in pairwise comparisons of software tools using LFQ intensities (p < 0.05).

Despite differences in absolute intensity distributions and differential expression outputs, FragPipe and MetaMorpheus showed broadly similar patterns in the number of differentially expressed proteins across extraction conditions. This suggests that relative fold-change trends may be partially conserved across software tools, although the magnitude and statistical significance of these changes remain sensitive to the underlying quantification and inference strategies. Consequently, while LFQ-based analyses can support comparative biological interpretation, the strength and stability of inferred differences depend on the choice of software pipeline. This distinction is particularly relevant for hair proteomics, where biomarker discovery often relies on relative abundance changes rather than absolute quantification.

## 1.5 Conclusion

Software choice influenced peptide identification, protein inference, functional annotation, and quantification in hair proteomics, even under harmonized search parameters and identical instrumentation. These findings highlight the importance of software selection for downstream biological interpretation and reproducibility in forensic and translational applications of hair analysis.

Substantial variability was observed in sequence coverage, PSMs, and protein group inference. This reflects differences in scoring strategies and peptide-to-protein mapping across software platforms. These differences also propagated into functional enrichment and physicochemical property profiles, indicating that software processing affects both proteome coverage and derived biological interpretation.

At the quantitative level, absolute LFQ intensities and differential expression outputs varied across tools. However, relative fold-change trends were partially conserved. FragPipe and MetaMorpheus showed broadly comparable differential expression patterns despite differences in intensity scaling, suggesting some consistency in directional interpretation, although magnitude and significance varied.

Interpretation of extraction-specific effects should be made with caution. DMC and SDD datasets were derived from different biological samples, whereas SDS and urea originated from the same source material. Observed differences likely reflect a combination of extraction chemistry, biological variability, and computational processing.

No single workflow consistently outperformed others across all metrics. This supports the use of multi-tool validation strategies in hair proteomics. Future work should incorporate a reference hair proteome to improve confidence in protein identification and enable standardized benchmarking across software platforms.

## References

1. Marsden, A.J., et al., Love is in the hair: arginine methylation of human hair proteins as novel cardiovascular biomarkers. Amino Acids, 2022. 54(4): p. 591–600.

2. Parker, G.J., et al., Demonstration of protein-based human identification using the hair shaft proteome. PLOS One, 2016. 11(9): p. 1–26.

3. Adeola, H.A., et al., No difference in the proteome of racially and geometrically classified scalp hair sample from a South African cohort: Preliminary findings. Journal of Proteomics, 2020. 226: p. 103892–103892.

4. Wu, J., et al., Deep coverage proteome analysis of hair shaft for forensic individual identification. Forensic Sci Int Genet, 2022. 60: p. 102742.

5. Florou, V.A., et al., Human hair as a diagnostic tool in medicine. Biochemistry and Biophysics Reports, 2025. 43: p. 102129.

6. Rogers, G.E., Hair follicle differentiation and regulation. The International Journal of Developmental Biology, 2004. 48(2-3): p. 163–170.

7. Buffoli, B., et al., The human hair: from anatomy to physiology. International Journal of Dermatology, 2014. 53(3): p. 331–341.

8. Robbins, C.R., Morphological, macromolecular structure and hair growth, in Chemical and Physical Behavior of Human Hair, C.R. Robbins, Editor. 2012, Springer-Verlag: Berlin. p. 1–104.

9. Wong, S.Y., et al., A high-yield two-hour protocol for extraction of human hair shaft proteins. PLOS One, 2016. 11(10): p. e0164993–e0164993.

10. Carlson, T.L., et al., Protein extraction from human anagen head hairs one millimeter or less in total length. BioTechniques, 2018. 64(4): p. 170–176.

11. Adav, S.S., et al., Studies on the proteome of human hair - Identification of histones and deamidated keratins. Nature Scientific Reports, 2018. 8(1): p. 1599–1610.

12. Bryson, W.G., et al., Improved two-dimensional electrophoretic mapping of Japanese human hair proteins; application to curved and straight Japanese human hairs; and protein identification by MALDI MS and MS/MS quadrupole time-of-flight mass spectrometry. 2020.

13. Barthélemy, N.R., et al., Proteomic tools for the investigation of human hair structural proteins and evidence of weakness sites on hair keratin coil segments. 2012. 421: p. 43–55.

14. Milan, J.A., et al., Comparison of protein expression levels and proteomically-inferred genotypes using human hair from different body sites. Forensic Science International: Genetics, 2019. 41: p. 19–23.

15. Laatsch, C.N., et al., Human hair shaft proteomic profiling: individual differences, site specificity and cuticle analysis. PeerJ, 2014. e506: p. 1–17.

16. Goecker, Z.C., et al., Optimal processing for proteomic genotyping of single human hairs. Forensic Science International: Genetics, 2020. 47: p. 102314.

17. Miller, R.M. and L.M. Smith, Overview and considerations in bottom-up proteomics. Analyst, 2023. 148(3): p. 475–486.

18. Dupree, E.J., et al., A critical review of bottom-up proteomics: The good, the bad, and the future of this field. Proteomes, 2020. 8(14): p. 1–26.

19. Chen, C., et al., Bioinformatics methods for mass spectrometry-based proteomics data analysis. International Journal of Molecular Sciences, 2020. 21(8).

20. Perkins, D.N., et al., Probability-based protein identification by searching sequence databases using mass spectrometry data. Electrophoresis, 1999. 20(18): p. 3551–3567.

21. Cox, J., et al., A practical guide to the MaxQuant computational platform for SILAC-based quantitative proteomics. Nature Protocols, 2009. 4(5): p. 698–705.

22. Tabb, D.L., J.K. Eng, and J.R. Yates, Protein identification by SEQUEST, in Proteome Research: Mass Spectrometry, P. James, Editor. 2001, Springer: Heidelberg. p. 125–142.

23. Bjornson, R.D., et al., X!!Tandem, an improved method for running X!Tandem in parallel on collections of commodity computers. Journal of Proteome Research, 2008. 7(1): p. 293–299.

24. Nesvizhskii, A.I., A survey of computational methods and error rate estimation procedures for peptide and protein identification in shotgun proteomics. Journal of Proteomics, 2010. 73(11): p. 2092–2123.

25. Bu, F., et al., Expression Profile of GINS Complex Predicts the Prognosis of Pancreatic Cancer Patients. Onco Targets Ther, 2020. 13: p. 11433–11444.

26. Nahnsen, S., et al., Tools for label-free peptide quantification. Molecular and Cellular Proteomics, 2013. 12(3): p. 549–556.

27. Sinitcyn, P., J.D. Rudolph, and J. Cox, Computational methods for understanding mass spectrometry–based shotgun proteomics data. Annual Review of Biomedical Data Science, 2018. 1(Volume 1, 2018): p. 207–234.

28. Adav, S.S., C.Y. Leung, and K.W. Ng, Profiling of hair proteome revealed individual demographics. Forensic Science International: Genetics, 2023. 66: p. 102914.

29. Chawade, A., et al., Data processing has major impact on the outcome of quantitative label-free LC-MS analysis. Journal of Proteome Research, 2015. 14(2): p. 676–87.

30. Fu, J., et al., Label-free proteome quantification and evaluation. Briefings in Bioinformatics, 2023. 24(1).

31. Griss, J., et al., Spectral Clustering Improves Label-Free Quantification of Low-Abundant Proteins. Journal of Proteome Research, 2019. 18(4): p. 1477–1485.

32. Kopczynski, D., A. Sickmann, and R. Ahrends, Computational proteomics tools for identification and quality control. Journal of Biotechnology, 2017. 261: p. 126–130.

33. Cox, J., et al., Accurate proteome-wide label-free quantification by delayed normalization and maximal peptide ratio extraction, termed MaxLFQ. Molecular and Cellular Proteomics, 2014. 13(9): p. 2513–26.

34. Cox, J. and M. Mann, MaxQuant enables high peptide identification rates, individualized p.p.b.-range mass accuracies and proteome-wide protein quantification. Nat Biotechnol, 2008. 26(12): p. 1367–72.

35. Kong, A.T., et al., MSFragger: ultrafast and comprehensive peptide identification in mass spectrometry-based proteomics. Nature Methods, 2017. 14(5): p. 513–520.

36. Yu, F., et al., Fast Quantitative Analysis of timsTOF PASEF Data with MSFragger and IonQuant. Molecular and Cellular Proteomics, 2020. 19(9): p. 1575–1585.

37. da Veiga Leprevost, F., et al., Philosopher: a versatile toolkit for shotgun proteomics data analysis. Nature Methods, 2020. 17(9): p. 869–870.

38. Solntsev, S.K., et al., Enhanced global post-translational modification discovery with MetaMorpheus. Journal of Proteome Research, 2018. 17(5): p. 1844–1851.

39. Miller, R.M., et al., Enhanced proteomic data analysis with MetaMorpheus, in Statistical Analysis of Proteomic Data: Methods and Tools, T. Burger, Editor. 2023, Springer US: New York, NY. p. 35–66.

40. Vaudel, M., et al., PeptideShaker enables reanalysis of MS-derived proteomics data sets. Nature Biotechnology, 2015. 33(1): p. 22–24.

41. Barsnes, H. and M. Vaudel, SearchGUI: A highly adaptable common interface for proteomics search and de novo engines. Journal of Proteome Research, 2018. 17(7): p. 2552–2555.

42. Oliveros, J.C. VENNY. An interactive tool for comparing lists with Venn Diagrams. 2007; Available from: http://bioinfogp.cnb.csic.es/tools/venny/index.html.

43. Szklarczyk, D., et al., STRING v11: protein–protein association networks with increased coverage, supporting functional discovery in genome-wide experimental datasets. Nucleic Acids Research, 2018. 47(D1): p. D607–D613.

44. Kim, S. and P.A. Pevzner, MS-GF+ makes progress towards a universal database search tool for proteomics. Nature Communications, 2014. 5.

45. Khalil, S. and M. Plisnier, Comparative analysis of MS/MS search algorithms in label-free shotgun proteomics for monitoring host-cell proteins using trapped ion mobility and ddaPASEF. Journal of Pharmaceutical and Biomedical Analysis Open, 2025. 6: p. 100082.

46. Yu, F., Y. Deng, and A.I. Nesvizhskii, MSFragger-DDA+ enhances peptide identification sensitivity with full isolation window search. Nature Communications, 2025. 16(1): p. 3329.

47. Craig, R. and R.C. Beavis, TANDEM: matching proteins with tandem mass spectra. Bioinformatics, 2004. 20(9): p. 1466–7.

48. Geer, L.Y., et al., Open mass spectrometry search algorithm. Journal of Proteome Research, 2004. 3(5): p. 958–964.

49. Miller, R.M., et al., Improved protein inference from multiple protease bottom-up mass spectrometry data. Journal of Proteome Research, 2019. 18(9): p. 3429–3438.

50. Zhang, B., M.C. Chambers, and D.L. Tabb, Proteomic parsimony through bipartite graph analysis improves accuracy and transparency. Journal of Proteome Research, 2007. 6(9): p. 3549–57.

51. Kapp, E.A., et al., An evaluation, comparison, and accurate benchmarking of several publicly available MS/MS search algorithms: sensitivity and specificity analysis. Proteomics, 2005. 5(13): p. 3475–3490.

52. Shteynberg, D., et al., Combining results of multiple search engines in proteomics. Mol Cell Proteomics, 2013. 12(9): p. 2383–93.

53. Audain, E., et al., In-depth analysis of protein inference algorithms using multiple search engines and well-defined metrics. Journal of Proteomics, 2017. 150: p. 170–182.

54. Peng, H., et al., Optimizing differential expression analysis for proteomics data via high-performing rules and ensemble inference. Nature Communications, 2024. 15(1): p. 3922.

55. Tabb, D.L., et al., Repeatability and reproducibility in proteomic identifications by liquid chromatography-tandem mass spectrometry. Journal of Proteome Research, 2010. 9(2): p. 761–76.

56. Lee, Y.J., R.H. Rice, and Y.M. Lee, Proteome analysis of human hair shaft: From protein identification to posttranslational modification. Molecular and Cellular Proteomics, 2006. 5(5): p. 789–800.

57. Solazzo, C., et al., Characterisation of novel α-keratin peptide markers for species identification in keratinous tissues using mass spectrometry. Rapid Communications in Mass Spectrometry, 2013. 27(23): p. 2685–2698.

58. Nie, S., et al., Maximizing hydrophobic peptide recovery in proteomics and antibody development using a mass spectrometry compatible surfactant. Analytical Biochemistry, 2022. 658: p. 114924.

59. Khamkar, V.S., M. Narasimhan, and R.B. Govekar, Improved identification of hydrophobic proteins by optimization of LC conditions within the LC-MS run-A practical strategy for scanty clinical samples. Biomedical Journal of Scientific & Technical Research, 2022. 43(4): p. 34886–34892.

60. Robbins, C.R., Chemical composition of different hair types, in Chemical and Physical Behavior of Human Hair. 2012, Springer-Verlag: Berlin. p. 105–176.

61. Plubell, D.L., et al., Putting Humpty Dumpty back together again: What does protein quantification mean in bottom-up proteomics? Journal of Proteome Research, 2022. 21(4): p. 891–898.

62. Zhao, Q., et al., 1-Dodecyl-3-methylimidazolium chloride-assisted sample preparation method for efficient integral membrane proteome analysis. Analytical Chemistry, 2014. 86(15): p. 7544–50.

63. Välikangas, T., T. Suomi, and L.L. Elo, A comprehensive evaluation of popular proteomics software workflows for label-free proteome quantification and imputation. Briefings in Bioinformatics, 2017. 19(6): p. 1344–1355.

