## Supplemental Tables 1 - 5 for "A comparison of four proteomics software for hair proteome analyses – A technical note"

Mukonyora, M.^1^

^1^Department of Medicine, University of Cape Town, Groote Schuur Hospital, Cape Town, South Africa.

***Supplementary 1: Summary of peptide identification and protein inference results***

| **Extraction method** | **Peptide identification and protein inference software** | **Validated peptide Spectrum Matches (PSMs)** | **Validated unique peptides** | **Proteins** | **Protein groups** | **Validated sequence coverage** |
| --- | --- | --- | --- | --- | --- | --- |
|  | SearchGUI (MSAmanda) & PeptideShaker | 61182 | 223 | 91 | 72 | 13.5% |
|  | SearchGUI (OMSSA) & PeptideShaker | 31449 | 169 | 47 | 45 | 15.6% |
|  | SearchGUI (MSGF) & PeptideShaker | 88299 | 817 | 164 | 140 | 17.8% |
| sodium dodecanoate | SearchGUI (XTandem) & PeptideShaker | 90521 | 702 | 183 | 147 | 15.6% |
|  | FragPipe | 11135 | 515 | 235 | 122 | 13.50% |
|  | MetaMorpheus | 16317 | 565 | 378 | 310 | 9% |
|  | MaxQuant | 15409 | 476 | 235 | 122 | 17% |
|  | SearchGUI (MSAmanda) & PeptideShaker | 7088 | 386 | 40 | 35 | 36.5% |
|  | SearchGUI (OMSSA) & PeptideShaker | 3188 | 321 | 49 | 47 | 35.4% |
|  | SearchGUI (MSGF) & PeptideShaker | 17130 | 666 | 78 | 62 | 38.6% |
| sodium dodecyl sulfate | SearchGUI (XTandem) & PeptideShaker | 16433 | 661 | 76 | 70 | 33.7% |
|  | FragPipe | 4259 | 389 | 141 | 93 | 34.0% |
|  | MetaMorpheus | 11075 | 418 | 155 | 109 | 15% |
|  | MaxQuant | 5653 | 443 | 178 | 89 | 27% |
|  | SearchGUI (MSAmanda) & PeptideShaker | 1296 | 193 | 60 | 41 | 24.5% |
|  | SearchGUI (OMSSA) & PeptideShaker | 562 | 77 | 45 | 44 | 15.8% |
|  | SearchGUI (MSGF) & PeptideShaker | 3336 | 396 | 90 | 73 | 28.1% |
| urea | SearchGUI (XTandem) & PeptideShaker | 3592 | 443 | 94 | 72 | 29.9% |
|  | FragPipe | 1006 | 517 | 116 | 83 | 29.0% |
|  | MetaMorpheus | 2823 | 192 | 115 | 80 | 19% |
|  | MaxQuant | 1342 | 289 | 148 | 80 | 23% |
|  | SearchGUI (MSAmanda) & PeptideShaker | 23006 | 1083 | 285 | 268 | 16.85% |
|  | SearchGUI (OMSSA) & PeptideShaker | 16206 | 548 | 260 | 256 | 17.4% |
|  | SearchGUI (MSGF) & PeptideShaker | 89737 | 4062 | 656 | 639 | 20.7% |
| dodecyl methylimidazolium chloride | SearchGUI (XTandem) & PeptideShaker | 142825 | 3034 | 537 | 528 | 18% |
|  | FragPipe | 21088 | 43985 | 1159 | 997 | 13.1% |
|  | MetaMorpheus | 289734 | 3334 | 1495 | 1382 | 11.20% |
|  | MaxQuant | 37658 | 2172 | 718 | 544 | 13.50% |

**Supplementary 2: Top 10 most variable proteins – DMC**

| **Gene** | **Tool** | **CV** |
| --- | --- | --- |
| *KRTAP3-2* | MaxQuant | 0.256718 |
| *KRTAP3-2* | FragPipe | 0.24991 |
| *TF* | MetaMorpheus | 0.230795 |
| *KRTAP3-2* | MetaMorpheus | 0.175575 |
| *LYZ* | MetaMorpheus | 0.175435 |
| *KRTAP1-1* | MaxQuant | 0.171373 |
| *UGDH* | MetaMorpheus | 0.159736 |
| *LOC100996750* | FragPipe | 0.154797 |
| *KRTAP1-1* | FragPipe | 0.150565 |
| *KRTAP4-7* | MaxQuant | 0.148158 |

**Supplementary 3: Top 10 most variable proteins – SDD**

| **Gene** | **Tool** | **CV** |
| --- | --- | --- |
| *VCP* | FragPipe | 0.153914 |
| *TXN* | MaxQuant | 0.145787 |
| *LAP3* | MetaMorpheus | 0.141758 |
| *LDHB* | FragPipe | 0.141679 |
| *FASN* | MaxQuant | 0.13415 |
| *KRTAP3-1* | FragPipe | 0.133505 |
| *ATP5F1B* | FragPipe | 0.122202 |
| *ALDH2* | MaxQuant | 0.117764 |
| *HSPA5* | FragPipe | 0.115811 |
| *KRTAP19-5* | FragPipe | 0.115135 |

**Supplementary 4: Top 10 most variable proteins – SDS**

| **Gene** | **Tool** | **CV** |
| --- | --- | --- |
| *H2BC12* | MetaMorpheus | 0.112156 |
| *KRTAP19-5* | MetaMorpheus | 0.098975 |
| *KRT84* | FragPipe | 0.060411 |
| *VSIG8* | FragPipe | 0.049858 |
| *KRTAP9-1* | MetaMorpheus | 0.046813 |
| *KRTAP4-1* | FragPipe | 0.0315 |
| *IGKC* | MetaMorpheus | 0.028781 |
| *KRTAP4-1* | MetaMorpheus | 0.027151 |
| GAPDHS | MaxQuant | 0.024926 |
| LOC100996750 | MetaMorpheus | 0.022304 |

**Supplementary 5: Top 10 most variable proteins – urea**

| Gene | Tool | CV |
| --- | --- | --- |
| KRTAP4-11 | MetaMorpheus | 0.107296 |
| KRTAP4-6 | FragPipe | 0.064416 |
| KRTAP7-1 | MetaMorpheus | 0.063585 |
| KRTAP2-2 | FragPipe | 0.062285 |
| KRT5 | MaxQuant | 0.035365 |
| IGHG3 | MaxQuant | 0.033031 |
| IGLL5 | FragPipe | 0.031877 |
| SFN | MaxQuant | 0.030178 |
| KRTAP9-7 | FragPipe | 0.02617 |
| KRT32 | MaxQuant | 0.025656 |
